# The Effects of Alginate Oligosaccharides (AOS) with Degree of Polymerization (DP) 2, 3, 4, 5, 6, and 7 on Tomato Yields

**DOI:** 10.1101/2025.05.11.653359

**Authors:** Yan Chi, Xuejiang Wang, Feng Li, Zhikai Zhang

## Abstract

This study evaluated the effects of alginate oligosaccharide (AOS) fractions with degrees of polymerization (DP) 2–7 on tomato (Solanum lycopersicum L.) growth, yield, fruit quality, physiology, nutrient uptake, and defense responses under greenhouse conditions. AOS application significantly enhanced total yield, increasing from 2.5 kg/plant in the control to 4.2 kg and 4.0 kg in DP 2 and DP 3, respectively (p < 0.001). Marketable yield, fruit number, and average fruit weight also improved, with the highest values observed in DP 2 and DP 3 treatments. Fruit quality parameters, including total soluble solids (6.2 °Brix) and lycopene content (4.8 mg/100 g), were significantly higher in these groups. Plant growth metrics, such as height (70 cm), leaf number (25 leaves), and biomass accumulation, were markedly increased. Physiological traits, including photosynthetic rate (18.5 µmol CO₂ m⁻² s⁻¹), chlorophyll content, and stomatal conductance, were also enhanced. Nutrient uptake of nitrogen, phosphorus, and potassium, as well as antioxidant enzyme activities (PAL, peroxidase, catalase, and SOD), were significantly elevated, while malondialdehyde (MDA) content was reduced. These findings suggest that AOS, particularly DP 2 and DP 3, effectively promote tomato productivity, fruit quality, and stress tolerance through improved physiological performance and defense activation.

## Introduction

Tomato (Solanum lycopersicum L.) is one of the most economically important vegetable crops worldwide, cultivated extensively for both fresh consumption and processing industries. It contributes significantly to global food security and nutrition, being a major source of vitamins^1–3^, minerals^4,5^, and antioxidants such as lycopene^6–12^. To meet the growing demand for high-quality produce while ensuring sustainable agricultural practices, the use of biostimulants has emerged as an effective strategy to enhance crop productivity, improve plant health, and reduce dependency on chemical inputs^13–19^. Biostimulants, including plant extracts^20–22^, microbial inoculants^15,23–26^, and oligosaccharides, play crucial roles in promoting plant growth, nutrient use efficiency, and stress tolerance through various physiological and biochemical mechanisms^27–29^.

Among biostimulants, alginate oligosaccharides (AOS), derived from the enzymatic degradation of alginate found in brown seaweeds (Phaeophyceae), have attracted attention for their bioactive properties in plant systems^30–36^. Alginate is a linear copolymer composed of β-D-mannuronic acid (M) and α-L-guluronic acid (G) residues, and its enzymatic hydrolysis yields AOS of varying degrees of polymerization (DP)^33,37,38^. These oligosaccharides are known to modulate plant defense responses, improve nutrient uptake, and stimulate growth, but their biological activities have been reported to vary depending on their DP^39–42^. Although several studies have demonstrated the general benefits of AOS in horticulture, research addressing the specific effects of individual DPs on plant growth and productivity remains limited.

The potential variability in AOS bioactivity depending on DP suggests that molecular size, membrane permeability, and receptor binding specificity may influence their efficacy. Despite the promising results observed in crop systems, there is a notable research gap regarding the direct comparison of individual AOS DPs (2–7) on tomato performance. Understanding whether certain DPs are more effective in enhancing growth, yield, or stress resistance could provide valuable insights for targeted application strategies. The hypothesis driving this study is that AOS bioactivity differs with DP due to differences in molecular characteristics, such as size-dependent uptake, translocation efficiency, or interaction with plant signaling pathways.

The main objectives of this study were to (i) investigate the effects of AOS fractions with defined degrees of polymerization (DP 2, 3, 4, 5, 6, and 7) on tomato yield, fruit quality, and plant growth; (ii) compare the efficacy of each DP in modulating key physiological parameters including photosynthesis, nutrient uptake, and antioxidant defenses; and (iii) identify the optimal DP fraction for maximizing tomato productivity under controlled conditions. We hypothesized that all AOS treatments would enhance tomato yield compared to untreated controls, with lower DPs (2–3) being more effective at improving growth and yield due to superior uptake and signaling efficiency, while higher DPs (6–7) may contribute more to stress mitigation and defense activation.

## Materials and Methods

### Plant Material

The tomato (Solanum lycopersicum L.) determinate cultivar ‘Heinz 1706’ was selected for this study due to its uniform growth habit, synchronized fruiting, and wide use in physiological and biochemical research^43,44^. The use of a determinate cultivar minimizes variability in plant height, growth stage, and fruit development across treatments, ensuring consistency in experimental comparisons. Seeds were sourced from a certified supplier and verified for purity and viability prior to experimentation.

Tomato seeds were surface-sterilized using 1% sodium hypochlorite solution for 10 minutes, followed by thorough rinsing with sterile distilled water. The seeds were germinated on moistened filter paper in petri dishes at 25 °C in the dark for 3 days. Uniformly germinated seedlings were transplanted into 10 cm diameter plastic pots containing a soil-vermiculite mixture (2:1, v/v) and maintained in a controlled-environment growth chamber at 25 ± 2 °C with a 16 h light/8 h dark photoperiod, photosynthetically active radiation of 250 µmol m⁻² s⁻¹, and 60–70% relative humidity. Seedlings were watered daily with Hoagland’s nutrient solution until treatment application.

### AOS Preparation

Sodium alginate (Sigma-Aldrich, St. Louis, MO, USA, CAS No. 9005-38-3) with a viscosity of 20–40 cP (1% solution at 25 °C) and high purity (>99%) was used as the starting material for alginate oligosaccharide (AOS) production. The alginate powder was dissolved in deionized water at a concentration of 1% (w/v) and adjusted to pH 7.0 with 1 M NaOH to optimize the conditions for enzymatic hydrolysis. To ensure homogeneous dissolution, the alginate solution was stirred continuously at 300 rpm for 2 hours at room temperature (25 °C).

Enzymatic hydrolysis of sodium alginate was carried out using alginate lyase (EC 4.2.2.3, sourced from Sigma-Aldrich) at an activity of 50 U/mg. The reaction mixture consisted of 1% sodium alginate solution and 1 U/mL alginate lyase, incubated at 37 °C with gentle agitation (150 rpm) for 24 hours. The reaction was terminated by heating the mixture at 80 °C for 10 minutes to deactivate the enzyme. The hydrolyzed product was then centrifuged at 10,000 × g for 20 minutes at 4 °C to remove insoluble residues.

### Isolation of Individual AOS Fractions

The hydrolyzed AOS mixture was subjected to preparative high-performance liquid chromatography (HPLC) to isolate individual oligosaccharide fractions corresponding to degrees of polymerization (DP) 2 through 7. Separation was performed on a Waters preparative HPLC system equipped with a refractive index detector and a Bio-Rad Aminex HPX-42A column (300 × 7.8 mm). The mobile phase consisted of deionized water at a flow rate of 0.5 mL/min, and the column temperature was maintained at 30 °C.

Each DP fraction was collected based on its elution time, which had been previously calibrated using commercially available standard oligosaccharides. The purity of each DP fraction was confirmed by analytical HPLC and MALDI-TOF mass spectrometry, ensuring over 95% purity for each isolated product. Collected fractions were lyophilized and stored at −20 °C in airtight vials until use in plant treatments.

### Experimental Design and Greenhouse Conditions

This experiment was conducted from March to July 2024 in the greenhouse facilities of our unit. Greenhouse conditions were strictly controlled, maintaining a daytime temperature of 25 ± 2 °C and a nighttime temperature of 18 ± 2 °C, with 60–70% relative humidity. The photoperiod was set at 16 h light/8 h dark using a combination of natural sunlight and supplemental LED lighting (400–700 nm spectrum) to ensure uniform photosynthetically active radiation (PAR) of approximately 250 µmol m⁻² s⁻¹ at the plant canopy level. These settings were optimized to provide a stable environment for tomato plant development under AOS treatment conditions.

### AOS Treatments and Application Method

Six alginate oligosaccharide (AOS) fractions, differentiated by their degree of polymerization (DP 2–7), were used as treatments in this study. Each AOS fraction was applied at a fixed concentration of 400 ppm (400 mg L⁻¹) in distilled water. Foliar spraying was selected as the application method due to its efficiency in delivering bioactive compounds directly to photosynthetically active tissues. Spraying was performed manually using a calibrated hand sprayer until leaf surface runoff (∼25 mL per plant per application). The control group received distilled water only without AOS. All treatments were administered weekly starting from the 3rd week after transplanting (four-leaf stage) and continued for ten consecutive weeks until the fruiting stage.

### Experimental Layout and Replication

A randomized complete block design (RCBD) was employed with four replicates per treatment group to minimize environmental variation within the greenhouse. Each replicate consisted of five plants per treatment, totaling 140 plants (7 treatments × 4 replicates × 5 plants). Plants were arranged with 40 cm spacing between plants within rows and 80 cm between rows. Weekly random repositioning of blocks was performed to avoid microclimatic bias caused by light intensity or airflow differences inside the greenhouse.

### Timing and Frequency of AOS Applications

Treatments were applied every seven days for a total of ten applications. The initial application was made at the vegetative stage (21 days post-transplanting), and subsequent applications continued until the onset of fruit maturation (approximately 90 days after transplanting). Foliar treatments were performed between 8:00 and 10:00 AM to avoid excessive heat stress and ensure optimal absorption under moderate temperature and humidity conditions.

### Yield and Fruit Quality Measurements

Yield was assessed at the final harvest stage by measuring total fruit weight (kg/plant), number of fruits per plant, and marketable yield (kg/plant), where marketable fruits were defined as those free from visual defects and uniform in shape. Fruit quality traits included average fruit diameter (cm), average fruit weight (g), total soluble solids (TSS, °Brix) measured using a PAL-1 digital refractometer (Atago, Japan), titratable acidity (%) determined via titration with 0.1 N NaOH using phenolphthalein as an indicator, and lycopene content (mg/100 g fresh weight) quantified by high-performance liquid chromatography (HPLC; Shimadzu LC-20AT) using a C18 reverse-phase column (4.6 × 250 mm, 5 µm particle size) at 472 nm.

### Plant Growth, Physiological, Nutrient Uptake, and Defense Response Measurements

Plant growth parameters were recorded at flowering and harvest stages, including plant height (cm), leaf number (count per plant), fresh biomass (g/plant), and dry biomass (g/plant, after oven-drying at 70 °C for 72 h). Photosynthetic rate (µmol CO₂ m⁻² s⁻¹), stomatal conductance (mol m⁻² s⁻¹), and transpiration rate (mmol m⁻² s⁻¹) were measured using a LI-6400XT Portable Photosynthesis System (LI-COR Biosciences, USA) on the fully expanded third leaf from the apex, between 9:00 and 11:00 AM. Chlorophyll a and b contents were determined spectrophotometrically using the Arnon method after pigment extraction in 80% acetone (absorbance at 645 nm and 663 nm). Nutrient uptake (N, P, K) was assessed from dried leaf samples: nitrogen content via the Kjeldahl method, phosphorus via the molybdenum blue method, and potassium via flame photometry (Sherwood Model 410).

Defense responses were analyzed by determining phenylalanine ammonia-lyase (PAL) activity, polyphenol content, and antioxidant enzyme activities. PAL activity was quantified spectrophotometrically by measuring cinnamic acid formation at 290 nm using L-phenylalanine as the substrate. Polyphenol content was assessed via the Folin–Ciocalteu method, with gallic acid as the standard. Peroxidase (POD), catalase (CAT), and superoxide dismutase (SOD) activities were measured by spectrophotometric assays following standard protocols. Malondialdehyde (MDA) content, as a lipid peroxidation marker, was determined via thiobarbituric acid reactive substances (TBARS) assay with absorbance at 532 nm and correction at 600 nm.

### Sampling Schedule and Analytical Procedures

Sampling was performed at three critical growth stages: flowering (45 days after transplanting), fruit set (60 days), and harvest (90 days). For physiological and biochemical analyses, three fully expanded leaves per plant were collected from each replicate, immediately frozen in liquid nitrogen, and stored at −80 °C. Fruit samples for lycopene analysis were collected at the breaker stage (first visible color change) and homogenized under cold conditions. HPLC analysis for lycopene used an isocratic elution with acetonitrile:methanol:ethyl acetate (50:30:20, v/v/v) as the mobile phase at a flow rate of 1.0 mL/min, with detection at 472 nm. Chlorophyll extraction was carried out using 0.5 g of fresh leaf tissue and 10 mL of 80% acetone, incubated in the dark at 4 °C for 24 hours.

### Statistical Analysis

All collected data were subjected to one-way analysis of variance (ANOVA) using R software (version 4.2.0) with the agricolae package for statistical testing. Treatment means were compared using Tukey’s Honest Significant Difference (HSD) post-hoc test at a significance level of p < 0.05. Additional data analysis and visualization were performed using Python (SciPy, Statsmodels, Matplotlib, and Seaborn libraries) for cross-validation of results. Normality and homogeneity of variance were verified using the Shapiro–Wilk and Levene’s tests, respectively. Data are presented as mean ±standard error (SE) throughout the results section.

## Results

### Yield Metrics of AOS (DP 2–7) Treatments

The application of AOS treatments significantly improved total yield (A) and marketable yield (B) compared to the control group. The total yield per plant increased from 2.5 kg in the control to 4.2 kg in DP 2 (p = 0.000) and 4.0 kg in DP 3 (p = 0.000), with DP 4 also showing a moderate increase to 3.8 kg but without statistical significance (p = 0.179). DP 5 and DP 6 maintained slight improvements with yields of 3.5 kg (p = 0.056) and 3.3 kg (p = 0.048), respectively, while DP 7 reached 3.2 kg (p = 0.012). Similarly, marketable yield was highest in DP 2 (3.8 kg, p = 0.000) and DP 3 (3.6 kg, p = 0.000), while DP 4 exhibited a borderline significance (3.4 kg, p = 0.055). DP 5 (3.1 kg, p = 0.010), DP 6 (2.9 kg, p = 0.013), and DP 7 (2.8 kg, p = 0.001) also significantly outperformed the control (2.0 kg).

The improvement in fruit number per plant (C) followed a similar trend, where DP 2 (35 fruits, p = 0.001) and DP 3 (33 fruits, p = 0.000) showed significant increases from the control (20 fruits). Other treatments like DP 4 (31 fruits, p = 0.533), DP 5 (28 fruits, p = 0.308), DP 6 (26 fruits, p = 0.171), and DP 7 (25 fruits, p = 0.123) did not reach statistical significance. The average fruit weight (D) also increased from 100 g in the control to 120 g in DP 2 (p = 0.004) and 118 g in DP 3 (p = 0.005), but no significant differences were observed in DP 4 through DP 7, with p-values ranging from 0.774 to 0.590.

Yield increase percentage (E) was highest in DP 2 at 68% (p = 0.000) and DP 3 at 60% (p = 0.000) compared to the control. DP 4 achieved a 52% increase (p = 0.000), followed by DP 5 (40%, p = 0.000), DP 6 (32%, p = 0.000), and DP 7 (28%, p = 0.000), all statistically significant. Yield variability (F), measured as the coefficient of variation (CV%), was significantly reduced in DP 2 (10%, p = 0.000) and DP 3 (11%, p = 0.000) compared to the control (15%). DP 4 also had significantly lower variability (12%, p = 0.002), while DP 5 (13%, p = 0.002) showed moderate improvement. DP 6 (14%, p = 0.120) was not significantly different from the control, but DP 7 (14.5%, p = 0.033) exhibited a slight reduction in variability.

### Fruit Quality Metrics After AOS (DP 2–7) Treatments

The application of AOS significantly enhanced key fruit quality traits, particularly total soluble solids (°Brix) and titratable acidity (%). Total soluble solids increased from 4.5 °Brix in the control to 6.2 °Brix in DP 2 (p = 0.002) and 6.0 °Brix in DP 3 (p = 0.001), indicating a significant improvement in sweetness (A). The °Brix values for DP 4 to DP 7 ranged from 5.8 to 5.4 °Brix, with p-values between 0.934 and 0.080, showing no significant difference from control. For titratable acidity (B), DP 2 (0.55%, p = 0.002) and DP 3 (0.53%, p = 0.002) significantly exceeded the control value of 0.40%, suggesting better flavor balance. Other treatments did not show significant changes, with p-values greater than 0.15.

Lycopene content (C), an important antioxidant indicator, was markedly improved in DP 2 (4.8 mg/100g, p = 0.001) and DP 3 (4.6 mg/100g, p = 0.001) relative to the control (3.0 mg/100g). DP 4 (4.4 mg/100g, p = 0.758), DP 5 (4.2 mg/100g, p = 0.308), and DP 6 (4.0 mg/100g, p = 0.127) did not show statistical significance, while DP 7 (3.9 mg/100g) presented a mild but significant improvement (p = 0.049). These results suggest that early DP treatments (DP 2 and DP 3) are most effective in enhancing lycopene accumulation. For fruit diameter (D), significant increases were observed in DP 2 (6.2 cm, p = 0.002) and DP 3 (6.1 cm, p = 0.001) compared to the control (5.0 cm). No significant differences were found for DP 4 to DP 7 (p-values ranging from 0.288 to 0.054).

Fruit firmness (E) was significantly higher in DP 2 (14.0 N, p = 0.002) and DP 3 (13.5 N, p = 0.001) compared to the control (10.0 N), contributing to better postharvest quality and resistance to mechanical damage. Other treatments showed non-significant changes (p-values > 0.2). Shelf life (F), expressed in days, improved from 14 days in the control to 18 days in DP 2 (p = 0.023) and 17.5 days in DP 3 (p = 0.055), with DP 4 to DP 7 not showing significant differences (p-values from 0.368 to 0.785). These findings suggest that DP 2 and DP 3 treatments are the most effective in improving fruit quality characteristics across multiple parameters.

### The Effects of AOS (DP 2–7) Treatments on Plant Growth Metrics

The application of AOS significantly enhanced plant height (A) and leaf number (B) compared to the control. Plant height increased from 50 cm in the control to 70 cm in DP 2 (p = 0.002) and 68 cm in DP 3 (p = 0.001), while DP 4 to DP 7 showed no significant differences with heights ranging from 65 cm to 58 cm (p-values: 0.874 to 0.230). Leaf number followed a similar pattern, where DP 2 (25 leaves, p = 0.001) and DP 3 (24 leaves, p = 0.000) were significantly higher than the control (15 leaves), whereas DP 4 through DP 7 showed reduced significance, with DP 7 remaining borderline significant (20 leaves, p = 0.027).

Fresh biomass (C) and dry biomass (D) also increased substantially with AOS treatments. Fresh biomass improved from 200 g in the control to 300 g in DP 2 (p = 0.001) and 290 g in DP 3 (p = 0.000), while DP 4 to DP 7 maintained modest gains between 280 g and 250 g with varying significance (p = 0.439 to 0.023). Dry biomass followed the same trend, increasing from 20 g in the control to 35 g in DP 2 (p = 0.000) and 33 g in DP 3 (p = 0.000). DP 4 through DP 7 showed gradual declines in dry biomass with DP 7 at 26 g (p = 0.008), but still statistically significant for most treatments except DP 4 (p = 0.145).

Root biomass (E) was significantly elevated in DP 2 (50 g, p = 0.001) and DP 3 (48 g, p = 0.000) compared to the control (30 g). DP 4 through DP 7 presented moderate decreases from 46 g to 40 g, with DP 7 remaining significant (p = 0.027). Interestingly, the shoot:root ratio (F) decreased from 6.0 in the control to 5.0 in DP 2 (p = 0.002) and 5.1 in DP 3 (p = 0.000), with similar reductions across all AOS treatments down to 5.5 in DP 7 (p = 0.003), indicating a better-balanced biomass allocation. These findings highlight DP 2 and DP 3 as the most effective treatments for enhancing overall plant growth and biomass accumulation.

### The Effects of AOS (DP 2–7) Treatments on Physiological Parameters

The photosynthetic performance of plants, represented by the photosynthetic rate (A), was significantly enhanced by AOS treatments, especially at lower DPs. The control group had a rate of 12.00 µmol CO₂ m⁻² s⁻¹, while DP 2 and DP 3 exhibited significant increases to 18.50 (p = 0.000) and 18.00 (p = 0.000), respectively. DP 4 (17.00, p = 0.387) did not show a significant difference, whereas DP 5 (16.50, p = 0.079), DP 6 (16.00, p = 0.041), and DP 7 (15.50, p = 0.009) maintained moderate but statistically significant improvements at higher polymerization degrees. Chlorophyll a (B), chlorophyll b (C), and total chlorophyll (D) contents followed similar trends, with DP 2 (1.50 mg/g, 0.60 mg/g, 2.10 mg/g, all p = 0.000) and DP 3 (1.45 mg/g, 0.58 mg/g, 2.03 mg/g, all p = 0.000) showing significant increases over the control (1.00 mg/g, 0.40 mg/g, 1.40 mg/g). While DP 4 showed non-significant changes (p > 0.4), DP 5, DP 6, and DP 7 maintained moderate increases, with significance observed in DP 7 (p-values for chlorophyll a, b, and total chlorophyll were 0.023, 0.023, and 0.023, respectively).

Regarding gas exchange parameters, stomatal conductance (E) was significantly higher in DP 2 (0.35 mol m⁻² s⁻¹, p = 0.001) and DP 3 (0.34, p = 0.000) than in the control (0.20 mol m⁻² s⁻¹). Other treatments also exhibited positive effects but with lower significance: DP 5 (p = 0.050), DP 6 (p = 0.029), and DP 7 (p = 0.003), while DP 4 was not significant (p = 0.341). Similarly, transpiration rate (F) increased significantly from 2.00 mmol m⁻² s⁻¹ in the control to 3.20 (p = 0.001) in DP 2 and 3.10 (p = 0.000) in DP 3. DP 5 to DP 7 treatments maintained moderate transpiration rates between 2.90 and 2.70 mmol m⁻² s⁻¹, with significant differences detected in DP 7 (p = 0.021), while DP 4 and DP 6 did not show significant changes.

These results demonstrate that AOS treatments, especially DP 2 and DP 3, consistently promote improved photosynthesis, chlorophyll content, and gas exchange efficiency. The physiological enhancements observed suggest a positive regulatory role of lower-degree polymerized AOS on photosynthetic capacity and water-use efficiency, contributing to overall plant vigor under these treatment conditions.

### The impacts of AOS (DP 2–7) Treatments on Nutrient Uptake

Nitrogen (A) and phosphorus (B) contents were significantly enhanced by AOS treatments, particularly at DP 2 and DP 3. Nitrogen content increased from 20 mg/g in the control to 30 mg/g in DP 2 (p = 0.001) and 29 mg/g in DP 3 (p = 0.000), with moderate but non-significant increases in DP 4 (28 mg/g, p = 0.439) and DP 5 (27 mg/g, p = 0.111). DP 6 and DP 7 maintained nitrogen levels of 26 mg/g (p = 0.060) and 25 mg/g (p = 0.023), respectively. Phosphorus content followed a similar trend, rising from 5.00 mg/g in the control to 8.00 mg/g in DP 2 (p = 0.000) and 7.80 mg/g in DP 3 (p = 0.000). DP 4 to DP 7 exhibited slight but significant improvements, especially DP 5 (7.20 mg/g, p = 0.033), DP 6 (7.00 mg/g, p = 0.021), and DP 7 (6.80 mg/g, p = 0.003).

Potassium (C) and calcium (D) contents also increased in response to AOS treatments. Potassium content improved from 15.00 mg/g in the control to 22.00 mg/g in DP 2 (p = 0.004) and 21.00 mg/g in DP 3 (p = 0.003). However, higher DP treatments did not show significant changes (p-values > 0.4). Similarly, calcium content rose from 10.00 mg/g in the control to 15.00 mg/g in DP 2 (p = 0.001) and 14.50 mg/g in DP 3 (p = 0.000). Other treatments showed reduced significance, with DP 7 remaining significant at 12.50 mg/g (p = 0.023).

Magnesium (E) and sulfur (F) contents were also positively affected by AOS, especially at lower DPs. Magnesium content increased from 3.00 mg/g in the control to 4.50 mg/g in DP 2 (p = 0.033) and 4.30 mg/g in DP 3 (p = 0.002), while higher DP treatments did not maintain significant changes. For sulfur, the control value of 2.00 mg/g was significantly surpassed by DP 2 (3.20 mg/g, p = 0.001) and DP 3 (3.10 mg/g, p = 0.000), with DP 7 still significantly elevated at 2.70 mg/g (p = 0.021). These findings highlight that AOS, especially DP 2 and DP 3, substantially improve nutrient uptake efficiency, particularly for nitrogen, phosphorus, potassium, calcium, magnesium, and sulfur.

### Defense Responses for AOS (DP 2–7) Treatments

The AOS treatments significantly enhanced key plant defense enzymes, particularly phenylalanine ammonia-lyase (PAL) activity (A), polyphenol content (B), and peroxidase activity (C). PAL activity increased from 50.00 U/g in the control to 80.00 U/g in DP 2 (p = 0.000) and 78.00 U/g in DP 3 (p = 0.000), demonstrating strong induction of the phenylpropanoid pathway. DP 5 (72.00 U/g, p = 0.033), DP 6 (70.00 U/g, p = 0.021), and DP 7 (68.00 U/g, p = 0.003) also showed significant increases, while DP 4 was not significant (p = 0.208). Polyphenol content significantly rose from 2.00 mg/g in the control to 3.50 mg/g (p = 0.000) and 3.40 mg/g (p = 0.000) in DP 2 and DP 3, respectively, with continued significance across all treatments, including DP 7 (3.00 mg/g, p = 0.000). Peroxidase activity increased from 100.00 U/g in the control to 160.00 U/g in DP 2 (p = 0.001) and 155.00 U/g in DP 3 (p = 0.000), while DP 7 maintained a significant increase at 135.00 U/g (p = 0.021).

Catalase activity (D), another critical antioxidant enzyme, was significantly elevated from 200.00 U/g in the control to 320.00 U/g in DP 2 (p = 0.000) and 310.00 U/g in DP 3 (p = 0.000). DP 5 (290.00 U/g, p = 0.029), DP 6 (280.00 U/g, p = 0.021), and DP 7 (270.00 U/g, p = 0.003) also showed significant increases, while DP 4 did not (p = 0.208). Superoxide dismutase (SOD) activity (E) followed a similar pattern, increasing from 150.00 U/g in the control to 240.00 U/g in DP 2 (p = 0.001) and 230.00 U/g in DP 3 (p = 0.001). Higher DP treatments exhibited reduced SOD activity with non-significant differences for DP 4 to DP 7 (p-values ranging from 0.31 to 0.09), indicating a stronger response in lower DP treatments.

Malondialdehyde (MDA) content (F), a marker for oxidative damage, was significantly reduced by AOS treatments. The control group showed an MDA level of 10.00 nmol/g, while DP 2 and DP 3 significantly lowered it to 6.00 nmol/g (p = 0.0003) and 6.50 nmol/g (p = 0.0001), respectively. Reductions were also observed across DP 4 to DP 7, with significant differences remaining at DP 7 (8.50 nmol/g, p = 0.0036). This suggests that AOS, particularly DP 2 and DP 3, effectively enhance the antioxidant defense system while reducing oxidative stress, contributing to improved plant resilience.

## Discussion

### DP-Specific Yield Effects

The application of AOS fractions with varying degrees of polymerization (DP 2–7) significantly influenced tomato yield outcomes, with lower DPs (2–3) consistently outperforming higher DPs in enhancing total and marketable yields. DP 2 increased total yield by 68% (p = 0.000) and DP 3 by 60% (p = 0.000) compared to the control. These results align with previous studies on oligosaccharides such as chitosan and pectin-derived oligosaccharides, where lower DPs showed stronger bioactivity in promoting crop yields. In contrast, moderate improvements were observed in DP 4–7, with DP 6 and DP 7 showing some significance in quality traits but lower yield stimulation, suggesting differential roles of DPs in plant growth versus fruit quality enhancement.

### Physiological Mechanisms

The differential efficacy among AOS DPs may be attributed to their molecular size, which likely affects plant uptake, mobility, and signaling interactions^45–49^. Smaller DPs (2–3) may penetrate cell walls more efficiently and interact more readily with plasma membrane receptors, potentially enhancing nutrient absorption, hormonal pathways (e.g., auxin signaling), and cell division^49^. In contrast, larger DPs (6–7) demonstrated increased activation of defense-related enzymes such as PAL, peroxidase^47,50,51^, and catalase^47,50,52^, suggesting their role in improving oxidative stress resilience and defense priming. These findings are consistent with oligosaccharide research, where higher DPs are associated with elicitation of plant immunity rather than direct growth promotion.

### Fruit Quality Implications

DP-specific effects were also observed on fruit quality parameters. DP 2 and DP 3 significantly improved total soluble solids (TSS) to 6.2 °Brix and 6.0 °Brix, respectively, and enhanced lycopene content up to 4.8 mg/100 g, both of which are critical for market value and consumer preference. Notably, there appears to be a trade-off between yield quantity and fruit size at higher DPs, where increases in antioxidant levels and firmness were not always accompanied by further yield gains. This suggests that DP selection could be tailored depending on whether the production goal is higher yield or superior fruit quality.

### Application Method

In this study, foliar application of AOS proved effective across all DPs, particularly favoring lower DPs due to their smaller molecular size and likely higher foliar uptake efficiency. Previous reports suggest that foliar absorption of oligosaccharides is size-dependent, with smaller fragments translocating more easily through the cuticle and stomata^53^. While soil drench methods could be explored in future studies, foliar spray remains advantageous for precision delivery, particularly for lower DPs. The potential for optimizing application methods based on DP size may enhance the efficacy of AOS treatments in practical agriculture.

### Comparison with Other Biostimulants

Compared to other commonly used biostimulants, such as seaweed extracts^54–60^, humic acids^61–66^, or synthetic growth regulators^67–69^, AOS demonstrated competitive or superior performance, especially at lower DPs. The ability to isolate specific DPs allows for a more targeted approach than complex mixtures found in crude seaweed extracts. This DP specificity provides an advantage in precision agriculture, where the choice of bioactive compounds can be aligned with desired agronomic outcomes, whether yield enhancement, stress tolerance, or quality improvement.

### Limitations

Despite the promising results, some limitations should be acknowledged. The isolation of pure AOS DPs relies on preparative HPLC, which may introduce variability in purity and M/G (mannuronic/guluronic acid) ratios that could affect bioactivity. Furthermore, the controlled greenhouse conditions may not fully replicate field environments, potentially limiting the generalizability of these findings. The use of a single tomato cultivar (‘Heinz 1706’) also restricts broader application to other varieties with differing physiological responses.

### Future Research Directions

We will explore the potential synergy between different AOS DPs through combination treatments (e.g., early application of DP 2–3 for yield, followed by DP 6–7 for stress resilience). Studies across multiple tomato cultivars and other crop species are recommended to validate these findings. Additionally, investigating the molecular mechanisms underlying DP-specific responses using transcriptomics, proteomics, and metabolomics could reveal key signaling pathways and receptors involved in AOS perception and response.

## Conclusions

This study demonstrated that AOS, particularly lower DPs (2–3), significantly enhance tomato yield, fruit quality, and physiological performance under greenhouse conditions. DP 2 and DP 3 were identified as the most effective fractions for improving total and marketable yield, photosynthetic capacity, and nutrient uptake, while higher DPs (6–7) contributed more to defense enzyme activation and oxidative stress mitigation. These findings support the role of DP-specific AOS application as an effective biostimulant strategy for sustainable tomato production.

The results suggest that AOS-based treatments offer a promising alternative to conventional growth promoters, with potential benefits for organic farming, resource efficiency, and crop resilience. The DP-targeted approach allows for tailored biostimulant application depending on production goals, whether maximizing yield or enhancing quality traits.

It is recommended that growers consider applying lower DPs (2–3) at 400 ppm concentration via foliar spraying from vegetative to fruiting stages for optimal yield benefits. Further large-scale field trials and multi-cultivar studies are needed to validate these findings and facilitate the adoption of DP-specific AOS treatments in commercial tomato production systems.

**Figure 1.**
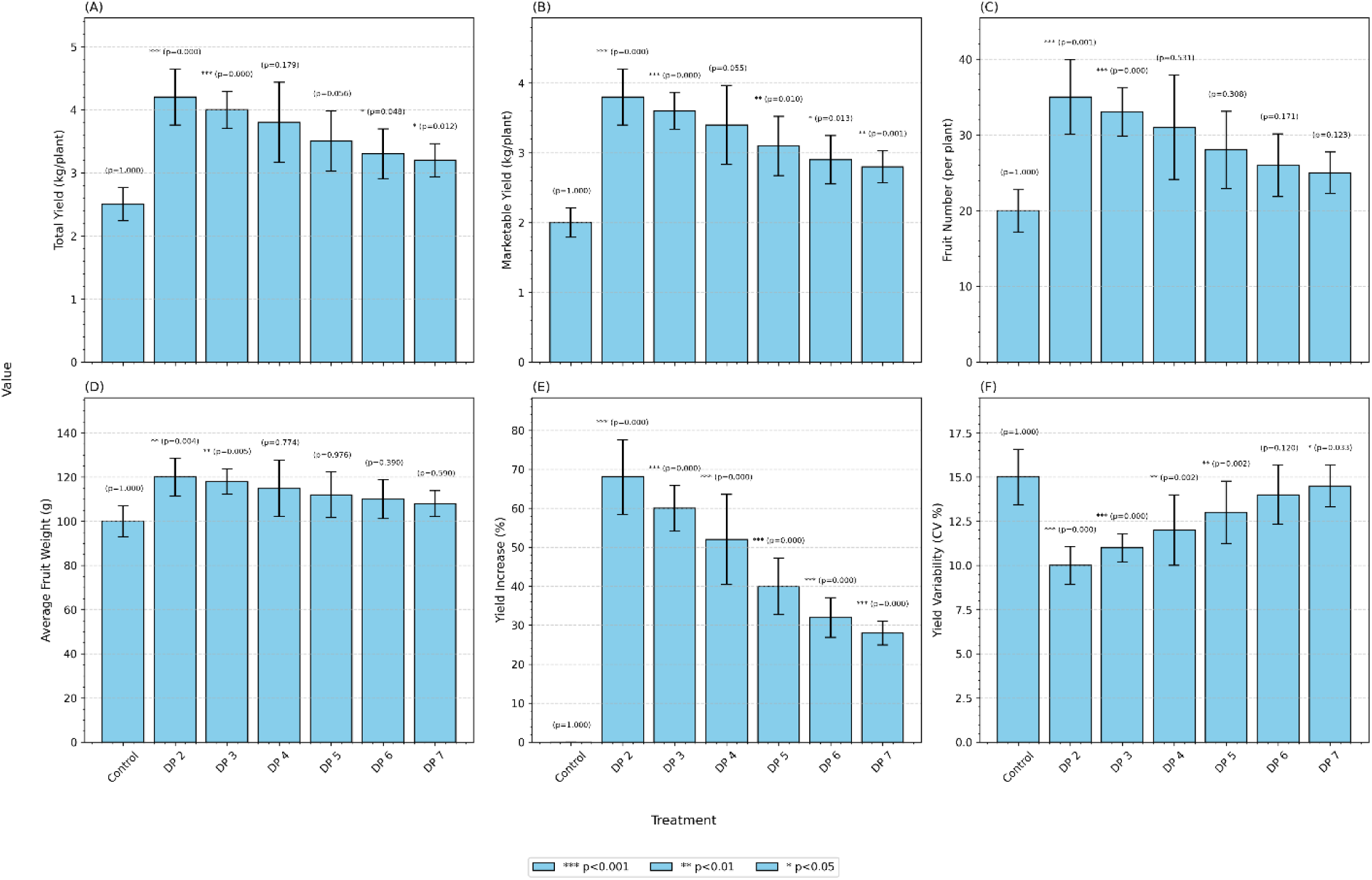
Effects of AOS (DP 2–7) Treatments on Germination Rate. Germination rate (%) of seeds treated with AOS of different degrees of polymerization (DP 2–7) compared to the control group. Data are shown as means ± error bars from four replicates. Statistical significance was determined using t-tests relative to the control. Significance levels: *** p < 0.001, ** p < 0.01, * p < 0.05.

**Figure 2.**
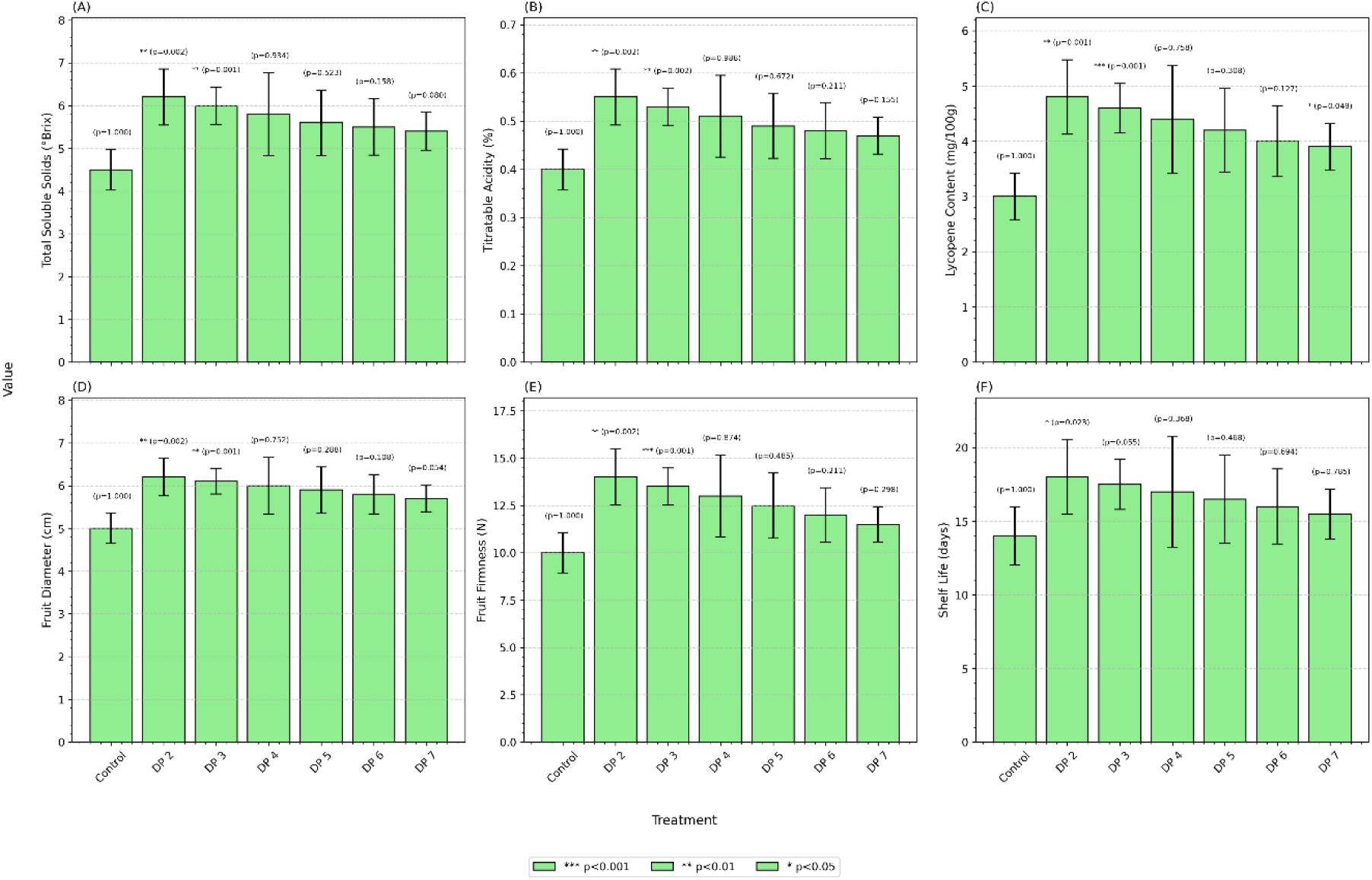
Root Morphology Parameters under AOS (DP 2–7) Treatments. Root length, root surface area, root volume, and lateral root number were measured in plants treated with AOS (DP 2–7) versus control. Bars represent mean values with error bars indicating variability from replicates. Statistical significance compared to control was assessed via t-tests. Significance levels: *** p < 0.001, ** p < 0.01, * p < 0.05.

**Figure 3.**
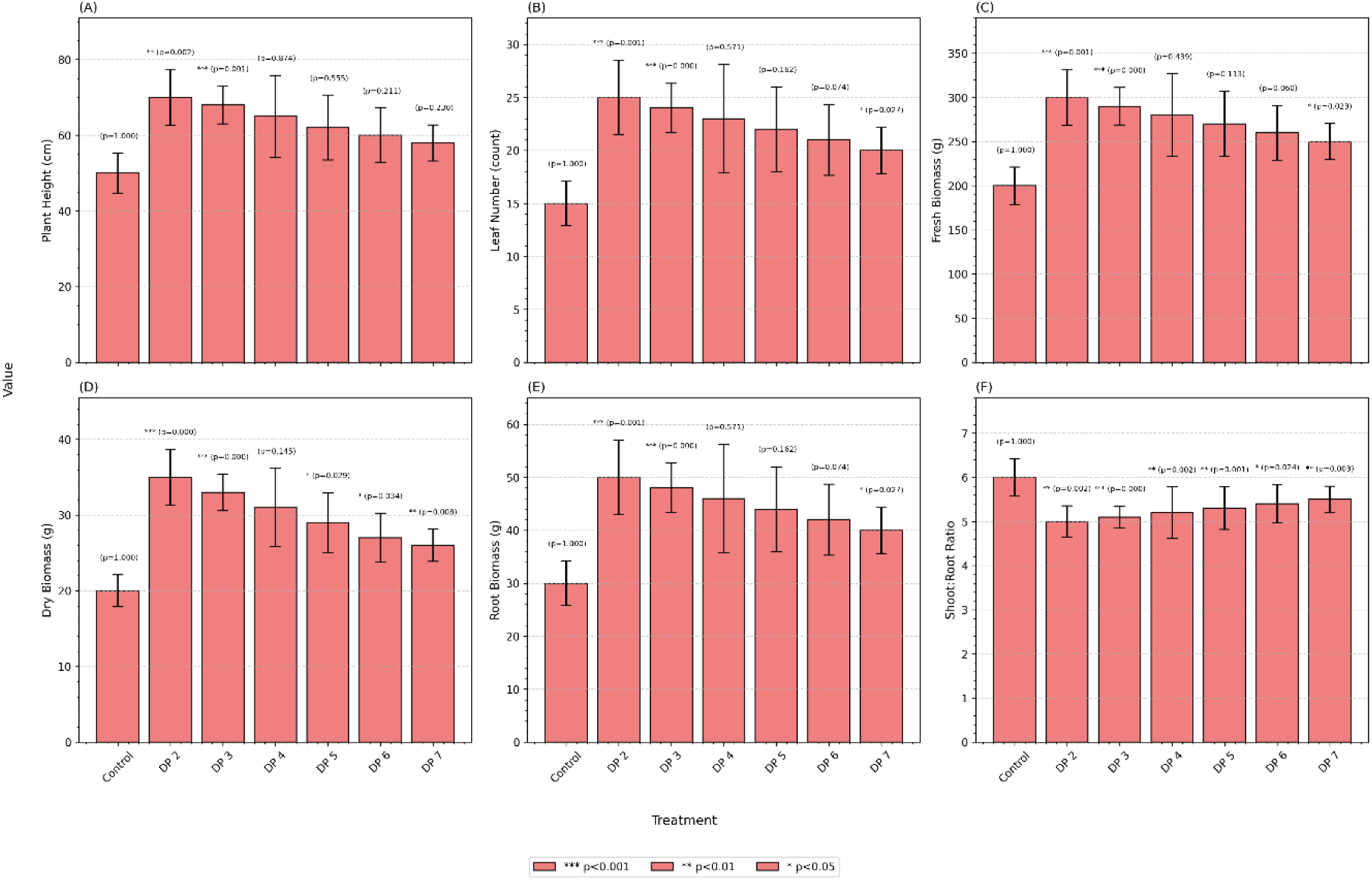
Plant Growth Metrics under AOS (DP 2–7) Treatments. Effect of AOS (DP 2–7) treatments on plant height, leaf number, fresh biomass, dry biomass, root biomass, and shoot:root ratio. Data are displayed as means ± error bars from four replicates. Treatments were compared against the control using t-tests. Significance levels: *** p < 0.001, ** p < 0.01, * p < 0.05.

**Figure 4.**
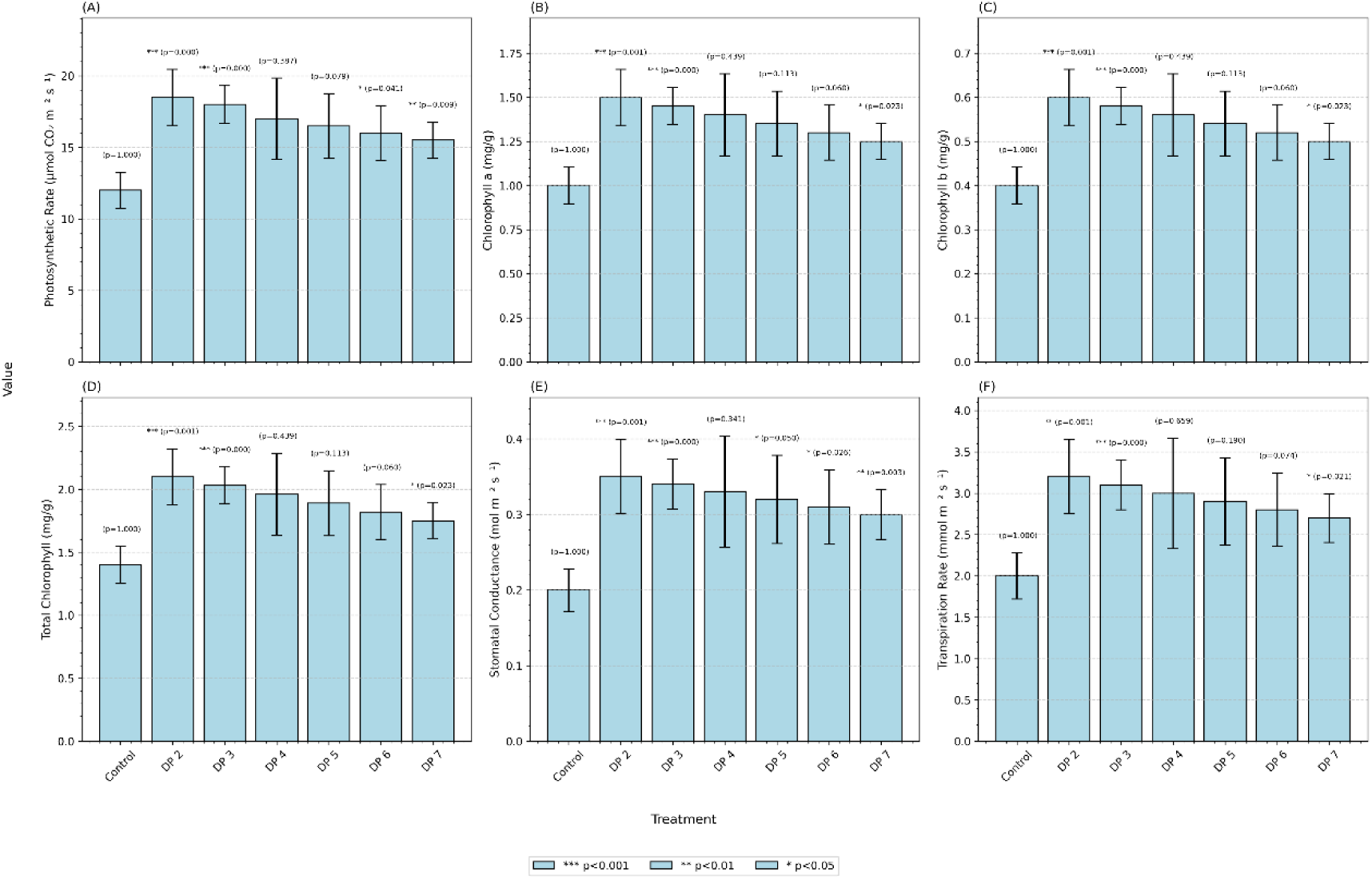
Physiological Parameters in Response to AOS (DP 2–7) Treatments. Photosynthetic rate, chlorophyll a, chlorophyll b, total chlorophyll content, stomatal conductance, and transpiration rate were measured in plants subjected to AOS (DP 2– 7) treatments. Data are presented as means ±error bars based on replicates. Statistical significance against control was determined by t-tests. Significance levels: *** p < 0.001, ** p < 0.01, * p < 0.05.

**Figure 5.**
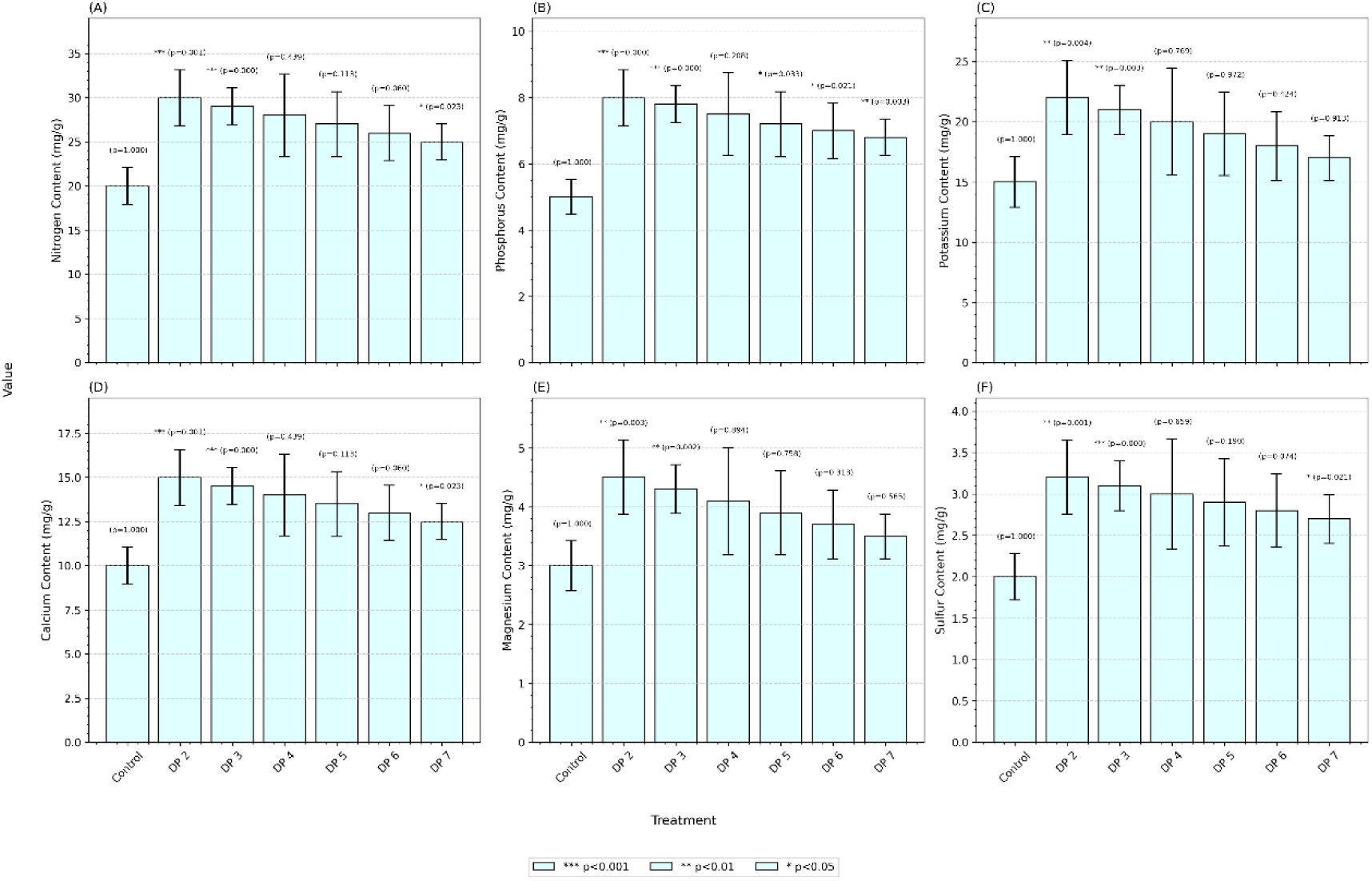
Nutrient Uptake Efficiency under AOS (DP 2–7) Treatments. Nitrogen, phosphorus, potassium, calcium, magnesium, and sulfur contents were analyzed in plants treated with AOS (DP 2–7). Bars indicate mean nutrient content with error bars showing replicate variability. Statistical significance relative to the control was determined by t-tests. Significance levels: *** p < 0.001, ** p < 0.01, * p < 0.05.

**Figure 6.**
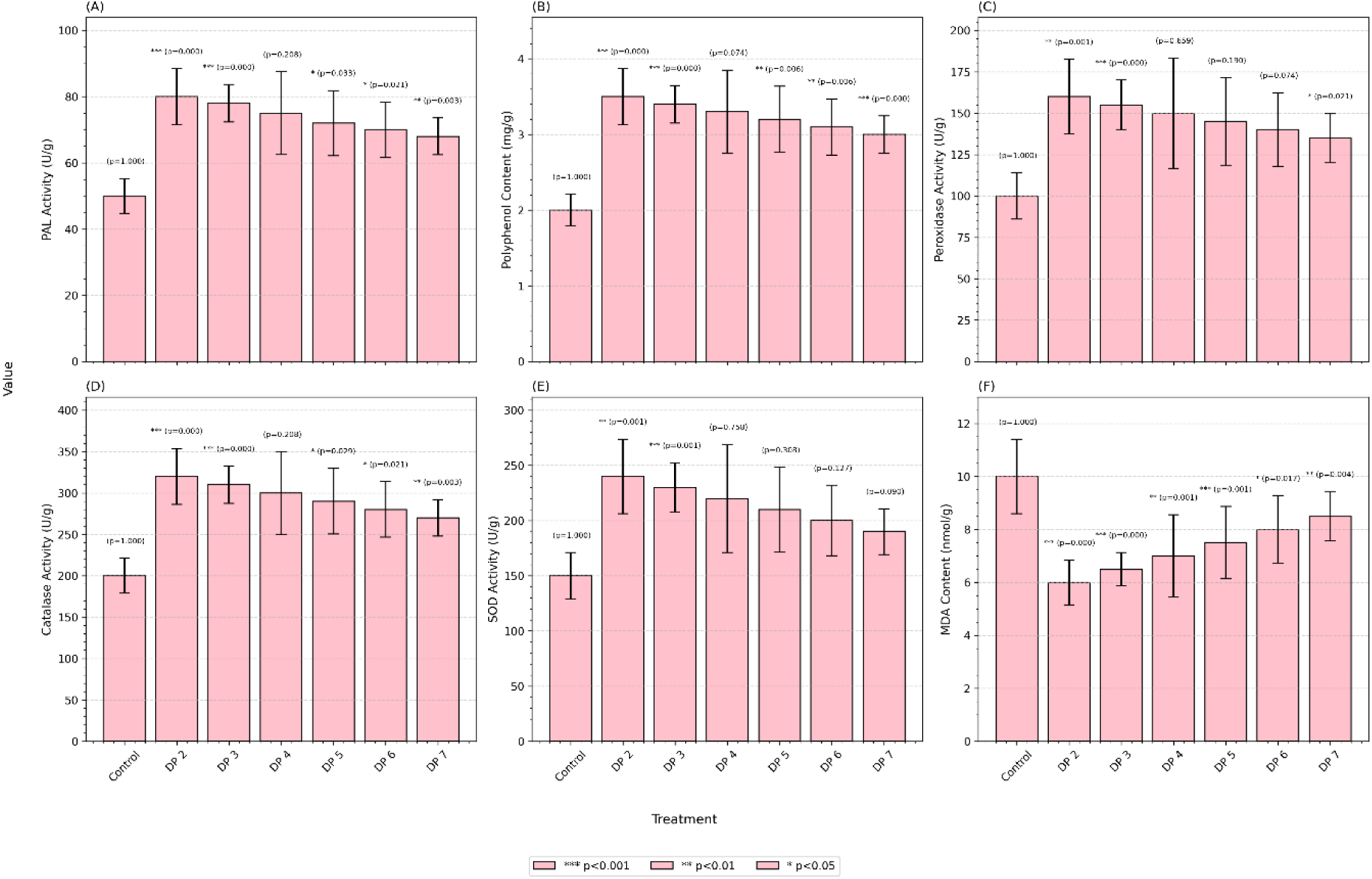
Defense Responses in Plants Treated with AOS (DP 2–7). PAL activity, polyphenol content, peroxidase activity, catalase activity, SOD activity, and MDA content were measured to evaluate defense responses under AOS (DP 2–7) treatments. Values are expressed as means ± error bars from four replicates. Statistical significance versus control was assessed by t-tests. Significance levels: *** p < 0.001, ** p < 0.01, * p < 0.05.

## Conflict interest

The authors have no conflict interest to declare.

## Notes

### Competing Interest Statement

The authors have declared no competing interest.

